# A Combination Adjuvant for the Induction of Potent Antiviral Immune Responses for a Recombinant SARS-CoV-2 Protein Vaccine

**DOI:** 10.1101/2021.02.18.431484

**Authors:** Sonia Jangra, Jeffrey J. Landers, Raveen Rathnasinghe, Jessica J. O’Konek, Katarzyna W. Janczak, Marilia Cascalho, Andrew A. Kennedy, Andrew W. Tai, James R. Baker, Michael Schotsaert, Pamela T. Wong

## Abstract

Several SARS-CoV-2 vaccines have received EUAs, but many issues remain unresolved, including duration of conferred immunity and breadth of cross-protection. Adjuvants that enhance and shape adaptive immune responses that confer broad protection against SARS-CoV-2 variants will be pivotal for long-term protection. We developed an intranasal, rationally designed adjuvant integrating a nanoemulsion (NE) that activates TLRs and NLRP3 with an RNA agonist of RIG-I (IVT DI). The combination adjuvant with spike protein antigen elicited robust responses to SARS-CoV-2 in mice, with markedly enhanced T_H_1-biased cellular responses and high virus-neutralizing antibody titers towards both homologous SARS-CoV-2 and a variant harboring the N501Y mutation shared by B1.1.7, B.1.351 and P.1 variants. Furthermore, passive transfer of vaccination-induced antibodies protected naive mice against heterologous viral challenge. NE/IVT DI enables mucosal vaccination, and has the potential to improve the immune profile of a variety of SARS-CoV-2 vaccine candidates to provide effective cross-protection against future drift variants.

## Introduction

The rapid spread of severe acute respiratory syndrome coronavirus 2 (SARS-CoV-2) has had a devastating impact on human health, with >145 million global cases of infection, and >3 million fatalities resulting since the start of the pandemic^1^. Several vaccines have received emergency use authorizations (EUAs) or are in late stage clinical testing^2–4^. However, many issues remain with these first generation vaccines, such as the duration and breadth of conferred immunity, whether vaccine induced immunity prevents transmission, and efficacy in cohorts with traditionally low response rates to vaccination such as the elderly and immunocompromised. The emergence of variants with higher transmissibility, increased virulence, and the potential for escape from current vaccines have instigated a new surge of infections, making it clear that successful control of the pandemic and mitigation of future outbreaks requires vaccines which provide not only robust and long-lasting protection, but also confer broad immunity towards current and future drift variants^5, 6^.

Multiple studies have confirmed that the potency of neutralizing antibodies (NAbs) induced by several of the vaccines currently deployed or in development (mRNA, adenovirus vectored, subunit) are impacted by different degrees to the current variants of concern (i.e. B.1.1.7, B.1.351, P.1)^7–10^. This reduction is especially prominent with the B.1.351 variant, which contains the N501Y mutation shared by the B.1.1.7 and P.1 variants, and two additional mutations (K417N, E484K) in the spike receptor binding domain (RBD)^11^. We and others have shown that the E484K substitution alone, significantly reduces the neutralization capacity of human convalescent and post-vaccination sera, which may leave people that have low NAb titers against current strains unprotected against newly emerging variants^11–14^. Therefore, improved strategies to induce higher titers of higher quality broadly neutralizing antibodies are needed as new variants continue to emerge. While NAbs directed towards the SARS-CoV-2 spike (S) protein are an important component of protective immunity, cellular immunity also plays a vital role in protection, especially in preventing severe disease and providing long-lasting immunity^15–17^. CD8^+^ T cell depletion partially abrogated protection from reinfection in NHPs^17^. Further, NAbs alone did not predict disease severity in COVID-19 patients, whereas both the presence of SARS-CoV-2 S-specific CD4^+^ and CD8^+^ T cells were significantly associated with less severe disease^15, 18–20^. Importantly, T cell epitopes are generally less susceptible to antigenic drift compared to antibody epitopes. Thus, a coordinated adaptive immune response composed of both robust NAbs and long-lasting CD4^+^ and CD8^+^ T cells is essential for strong protection, and will be critical to a successful vaccination strategy for broad protection against COVID-19.

Here, we demonstrate a rationally designed combination mucosal (intranasal (IN)) adjuvant that enhances the quality of the immune response to SARS-CoV-2 induced by an S protein subunit antigen. Adjuvants can facilitate induction of high levels of NAbs and robust protective T cell responses and help promote more durable immune memory. Furthermore, rational adjuvant design allows for shaping or skewing of vaccine responses towards effective, broader and more potent immunity against drifted viruses. Viruses that induce long-lasting immunity through natural infection typically stimulate strong innate immune responses through the activation of Toll-, RIG-I-, and NOD-like receptors (TLRs, RLRs, NLRs). However, SARS-CoV-2 and –CoV infections induce muted innate responses due to several immune-evasion tactics, including avoidance of RLR recognition and direct inhibition of RIG-I and downstream effector molecules, resulting in weaker induction of key cytokines including type-I interferons (IFN-Is) and poor activation of IFN-I associated pathways^21–23^. As IFN-Is are master activators of the antiviral defense program, and are essential to priming adaptive T cell responses and shaping effector and memory T cell pools, this weak innate response likely contributes to the large variability in magnitude and duration of immune responses observed in infected patients^24^. TLR signaling also drives T cell responses and affinity maturation of antiviral antibodies. As induction of appropriate innate responses is crucial for promoting durable, and quantitatively and qualitatively improved adaptive responses, we designed a combination adjuvant containing agonists for all three key innate receptor classes involved in SARS-CoV-2 immunity. The combined adjuvant integrates a nanoemulsion-based adjuvant (NE) that activates TLRs and NLRP3 with an RNA agonist of RIG-I (IVT DI). NE has established Phase I clinical safety as an IN adjuvant for influenza vaccines, and is currently being tested in another human study^25–27^. NE is composed of an oil-in-water emulsion of soybean oil, a nonionic (Tween80) and cationic (cetylpyridinium chloride) surfactant, and ethanol^28, 29^. NE-based immunity is both mucosal and systemic and has a pronounced T_H_1/T_H_17 bias, including an increase in antigen specific CD4^+^ and CD8^+^T cells^29–35^. Its activity is mediated at least in part through TLR2 and 4 activation, as well as through induction of immunogenic apoptosis which leads indirectly to NLRP3 activation^30, 35^. As an IN adjuvant, NE imparts robust protective immunity to viruses such as influenza virus, HSV-2, and RSV, without inducing clinical or histopathological signs of immune potentiation. IVT DI is an *in vitro* transcribed RNA consisting of the full-length (546nt) copy-back defective interfering RNA of Sendai virus strain Cantell^36, 37^. The hairpin structure of IVT DI, along with its dsRNA panhandle and 5’ triphosphate, make it a potent and selective RIG-I agonist, and thus, a strong inducer of IFN-Is and interferon-stimulated genes (ISGs).

We have previously shown that combining NE and IVT DI (NE/IVT) synergistically enhances protective immune responses towards influenza virus when administered IN, leading to improved antibody responses (with shortened kinetics, increased avidity, and viral neutralization) and broadened cross-subtype recognition, and induced a robust antigen specific cellular response with markedly magnified T_H_1 bias^38^. In these current studies, we immunized animals using this two-component adjuvant with the recombinant SARS-CoV-2 S1 subunit-a primary target for NAbs as it contains the RBD, which binds to the ACE2 receptor on target cells. We demonstrate that adjuvanting S1 with NE/IVT, markedly improves the magnitude and quality of the antibody responses towards both the homologous SARS-CoV-2 virus and a divergent mouse-adapted variant (MA-CoV2) harboring the N501Y substitution in the S protein found in the B.1.1.7, B.1.351, and P.1 variants. Passive transfer of vaccine-induced antibodies conferred robust protection against challenge with the heterologous SARS-CoV-2 variant, and resulted in sterilizing immunity in naïve mice. Moreover, robust antigen-specific cellular immune responses with a magnified T_H_1 bias along with a T_H_17 response were induced with NE/IVT. The combined adjuvant is compatible with both whole virus and recombinant protein antigens and thus provides a flexible platform that can improve the immune profile of several current and future SARS-CoV-2 vaccine candidates, and enable their use through the IN route, providing benefits unique to mucosal immunization.

## Results

### Safety and acute response evaluation

Extensive characterization of the NE adjuvant has been performed in multiple animal models as well as in humans, each demonstrating optimal safety profiles for use of the adjuvant through both intramuscular (IM) and IN routes^34, 39–41^. However, to examine whether multivalent stimulation of TLRs, NLRs, and RIG-I with the combined NE/IVT leads to over-activation of inflammatory responses, we evaluated the acute phase cytokine response by measuring levels of representative cytokines (IL-6, TNF-α, IL12p70, and IFN-γ) in the serum at 6 and 12 h post-immunization using recombinant SARS-CoV-2 RBD as a model antigen. Mice were immunized IN with 10 μg of RBD alone, or combined with NE (20% w/v) or NE/IVT (20% NE/0.5 μg IVT DI)—adjuvant doses used in all subsequent studies—in a total volume of 12 μL per mouse. Minimal or no acute inflammatory cytokines were detectable at these time points, with only IL-6 being modestly elevated in the NE/RBD (mean 34.4 pg/mL) and NE/IVT/RBD (mean 74.7 pg/mL) groups, as compared to the RBD only group (mean 11.6 pg/mL). No significant TNF-α, IL12p70, or IFN-γ were detectable. These results are consistent with the safety profile of the NE alone, supporting a lack of systemic toxicity with the combined adjuvant. These results are also consistent with the lack of significant changes in body temperature or weight in NE/IVT immunized mice in our previous studies with influenza virus antigens (data not shown).

### Characterization of the humoral response induced by immunization with SARS-CoV-2 S1

The impact of NE/IVT on the humoral response was assessed using recombinant SARS-CoV-2 S1 subunit as the antigen for immunization. The S1 contains the RBD which forms the necessary interactions with the human ACE2 receptor (hACE2) required for viral entry. We selected S1 for these initial studies rather than the RBD itself, as while the majority of key neutralizing epitopes map to the RBD, epitopes outside the RBD, particularly those proximal to it have also been found to contribute to neutralization. Moreover, full S1 provides additional antigenic sites outside of the RBD less prone to selective pressure and may thus impart improved crossvariant immunity. Mice were immunized IN with 15 μg of S1 alone (S1 only) or with 20% NE (NE/S1), or 20% NE/0.5 μg IVT DI (NE/IVT/S1) according to a prime/boost/boost schedule at a four-week interval. Serum S1-specific total IgG titers were measured two weeks after each immunization (Figure 2). Two weeks after the first immunization, antigen-specific IgG was not detectable in the S1 only group or in the majority of animals immunized with the adjuvants: NE/S1 and NE/IVT/S1 (Figure 2A). Each adjuvanted group had a single high responder with S1-specific IgG titers of 1:100, and 1:6250 for the NE/S1 and NE/IVT/S1 groups, respectively, already suggesting the potential for an improved synergistic effect in early antibody titers with the combined adjuvant upon further optimization of antigen dose. S1-specific IgG increased significantly in both adjuvanted groups after the second immunization, resulting in comparable geomean IgG titers (GMTs) in the range of ~10^5^ for the NE/S1 and NE/IVT/S1 groups, respectively (Figure 2B), which were further enhanced by the third immunization to titers of ~10^6^ (Figure 2C). In contrast, the S1 only group showed minimal IgG even after the second immunization, and reached only a titer of 1:200 after the third immunization.

**Figure 1.**
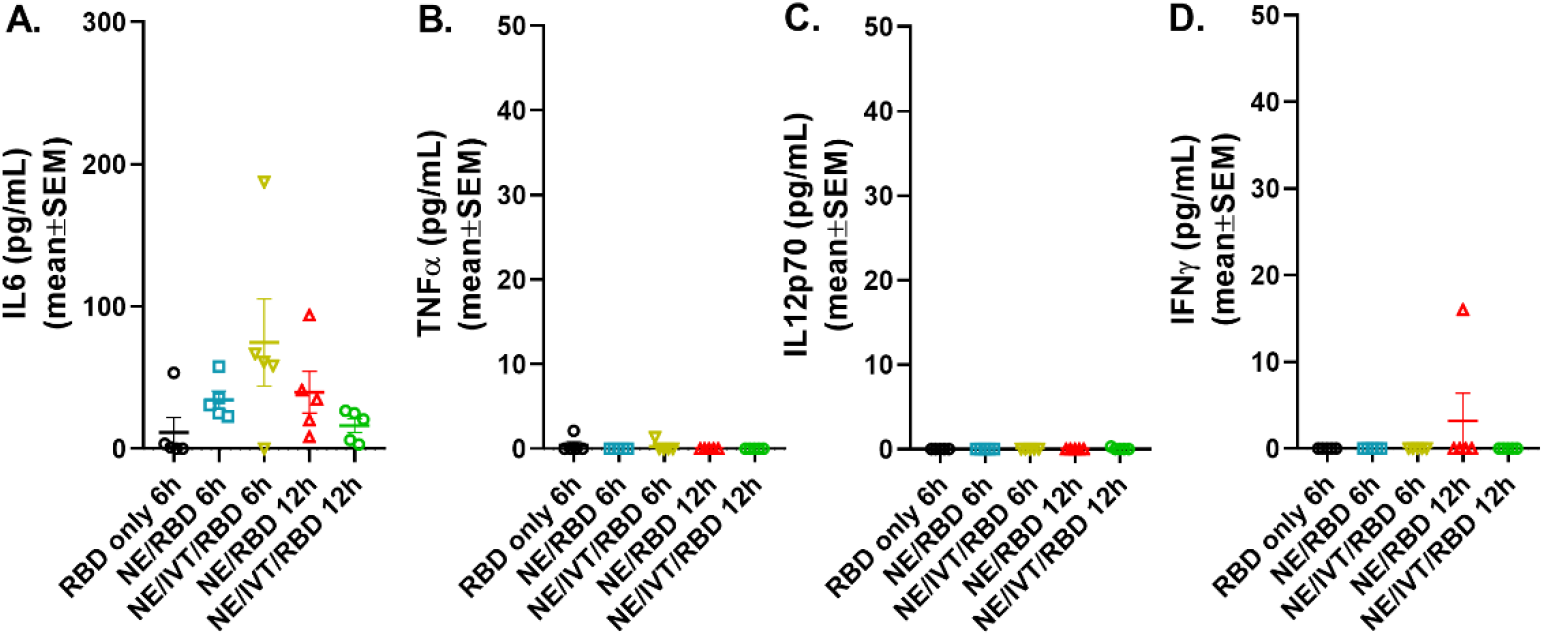
Acute phase cytokine response after immunization with NE and NE/IVT. Acute cytokines were assessed in the serum upon by measuring (A) IL-6, (B) TNF-α, (C) IL12p70, and (D) IFN-γ by multiplex immunoassay at 6 and 12 h post-IN immunization with 10 μg of RBD only, or with 20% NE, or 20% NE/0.5 μg IVT DI.

**Figure 2.**
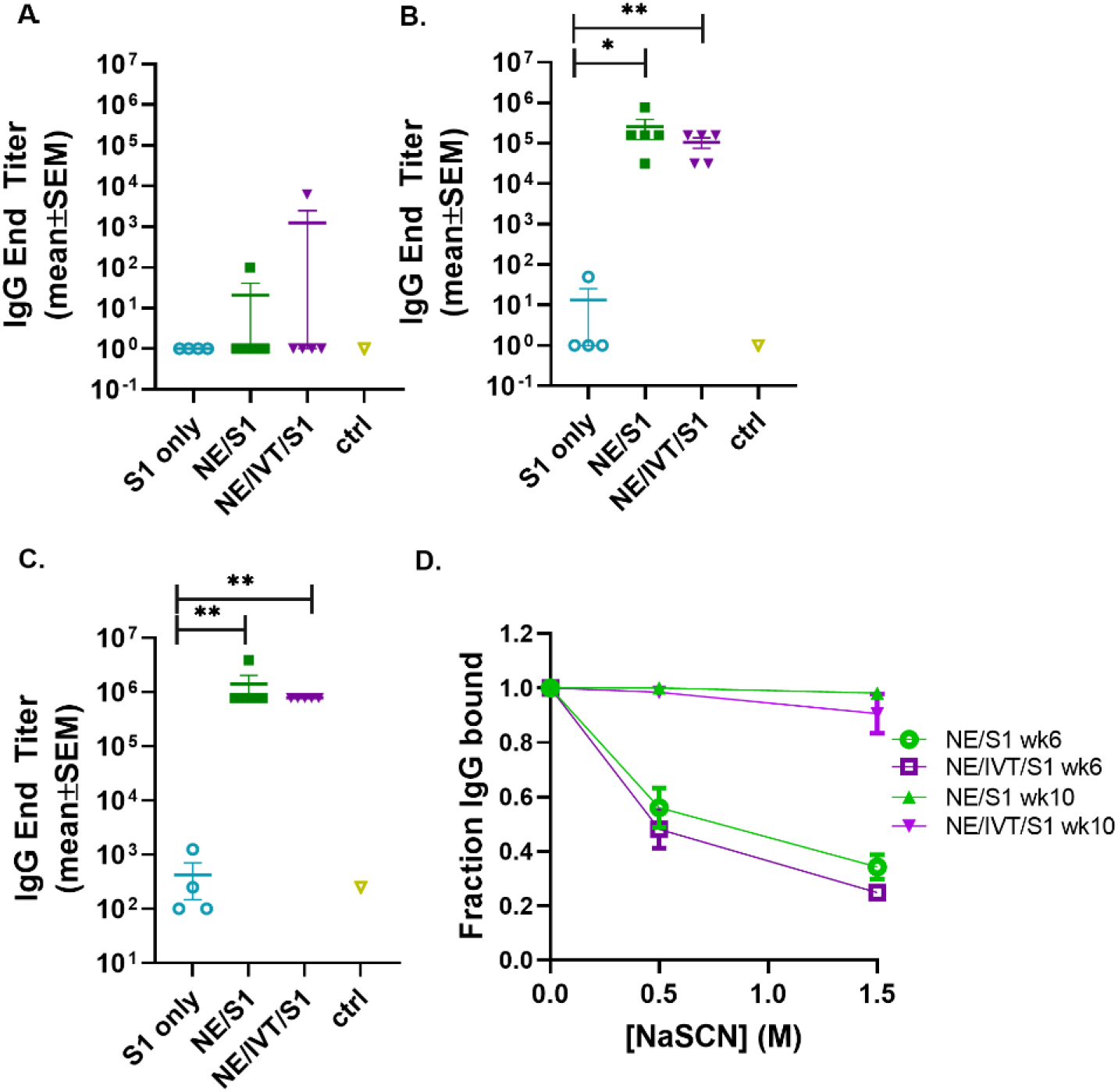
Serum S1-specific IgG titers induced by immunization with NE and NE/IVT and comparison of antibody avidity. Serum S1-specific IgG measured in mice immunized IN with 15 μg S1 alone, or with 20% NE, or 20%/0.5 μg IVT DI. S1-specific IgG was measured 2 weeks after each immunization at (A) 2 wks post-initial immunization (prime), (B) 6 wks post-initial immunization (prime/boost), and (C) 10 wks post-initial immunization (prime/boost/boost). (*p<0.05, **p<0.01 by Mann-Whitney U test). (D) Antibody avidity for S1-specific IgG measured by NaSCN elution for a 1:1,250 dilution of serum for the 6 wk and 10 wk sera. The control group (ctrl) consisted of untreated mice.

As we have previously shown that NE/IVT significantly enhances the avidity of antigen-specific IgG from mice immunized with whole inactivated influenza virus compared to the NE alone, we next examined whether the combined adjuvant also improved the avidity of the induced S1-specific antibodies by NaSCN chaotrope elution (Figure 2D)^38^. The NE/S1 and NE/IVT/S1 groups displayed identical antibody avidity for S1 after two immunizations (wk 6), which was significantly enhanced in both groups after the third immunization (wk 10). Both NE and NE/IVT induced exceptionally high affinity antibodies after three immunizations, with 95-100% of the S1-specific IgG remaining bound even upon elution with a high (1.5M) concentration of NaSCN at the highest dilution of serum tested (1:1,250). Such strong avidity was significantly greater than that of the antibodies evaluated from human convalescent sera, which had only 20-60% of the S1 antibodies remaining bound under a less stringent elution condition of 1 M NaSCN^42^.

To compare the IgG subclass distributions induced by NE and NE/IVT, the relative titers of IgG1, IgG2b, and IgG2c were measured for the wk 10 sera (Figure 3). Subclass analysis suggests that a balanced T_H_1/T_H_2 profile was elicited for both the NE and NE/IVT adjuvants, as has been observed in previous studies with other antigens ^33, 38, 43, 44^. Equivalent titers of IgG1 were induced by both NE and NE/IVT with GMTs of 5.7×10^5^ and 4.1×10^5^, respectively (Figure 3A).

**Figure 3.**
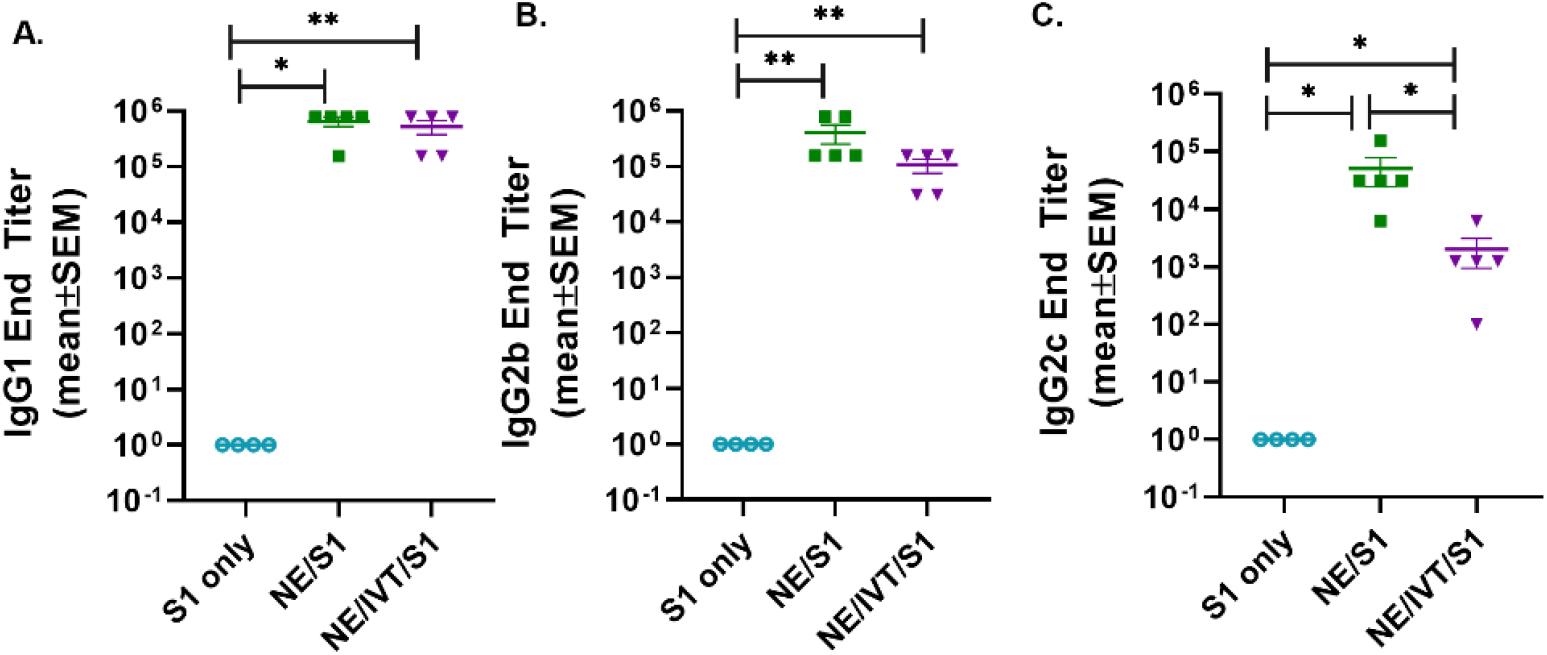
Serum S1-specific IgG subclass distributions induced by immunization. Levels of (A) IgG1, (B) IgG2b, and (C) IgG2c were measured in mice immunized IN with 15 μg S1 alone, or with 20% NE, or 20%/0.5 μg IVT DI after the last boost immunization, 10 wks post-initial immunization. (*p<0.05, **p<0.01 by Mann-Whitney U test).

High titers of T_H_1 associated subclasses were also observed for both adjuvants. S1-specific IgG2b GMTs of 3.0×10^5^ and 8.2×10^4^ (Figure 3B), and IgG2c GMTs of 3.1×10^4^ and 1.0×10^3^ were observed for NE and NE/IVT adjuvanted groups, respectively (Figure 3C). Interestingly, IgG2 subclasses were slightly reduced by the presence of IVT DI. Importantly, this balanced T_H_1/T_H_2 profile in combination with the cytokine data presented below suggest that these adjuvants avoid the strongly T_H_2-biased immune responses that have previously been linked to vaccine associated enhanced respiratory disease (VAERD) in SARS-CoV and RSV vaccine candidates adjuvanted with alum^45–49^.

To examine the functionality of the induced antibodies, viral neutralization was assessed using virus stock prepared from the clinical isolate, 2019-nCoV/USA-WA1/2020, which we classify as “WT” SARS-CoV-2. (Figure 4A, B). WT SARS-CoV-2 shares a high degree of homology with the Wuhan-Hu-1 isolate from which the S1 subunit used for immunization was derived. To examine the ability of the antibodies from the immunized mice to neutralize a variant with mutations in the S protein, we repeated the neutralization assay with our mouse-adapted SARS-CoV-2 virus (MA-SARS-CoV-2). The MA-virus was generated by serial passaging the WT virus isolate first in the lungs of immune compromised and then in immune competent mice of different backgrounds to optimize mouse virulence, as previously described^50^. As the WT virus is unable to use the endogenously expressed mouse ACE2 receptor (mACE2) for entry, productive infection of the murine respiratory tract is inefficient. Serial passaging selected for adaptive mutations which allowed the virus to optimally bind and use mACE2 for productive infection. The MA-SARS-CoV-2 S protein contains two amino acid (aa) mutations compared to the WT virus from which it was derived, including N501Y and H655Y, and a four aa insertion within the S1 subunit^50^. The N501Y substitution has previously been reported by other groups in an independent mouse adaptation of SARS-CoV-2, and is thus likely to be important for increasing affinity to the mACE2 receptor. Importantly, the N501Y mutation is shared by most of the recently identified variants of concern, B1.1.7, B.135.1, and P.1, among others, and is thought to play a role in the increased human to human transmissibility observed for these variants by increasing the affinity of the S protein for the hACE2 receptor^7^. In addition to the changes in the S protein, the MA-virus also contains three other mutations relative to the Wuhan-Hu-1 isolate: S194T in the nucleoprotein, T7I in the M protein, and L84S in ORF8. The L84S mutation is however present in the USA-WA1/2020 strain and is most likely not due to mouse adaptation.

**Figure 4.**
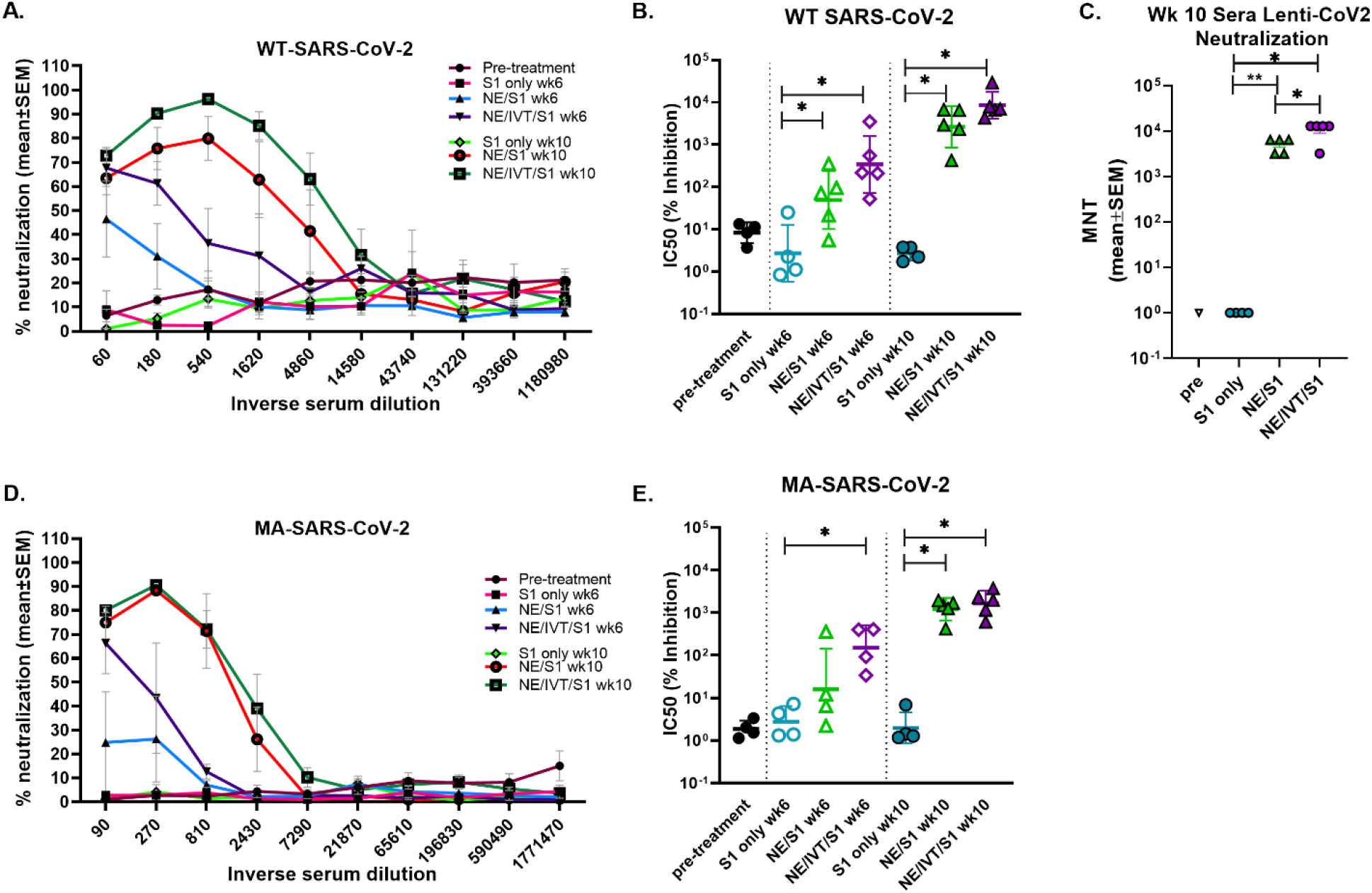
Virus neutralization titers to homologous and heterologous SARS-CoV-2 following immunization. Neutralizing antibody titers in serum from mice receiving two (wk 6) or three (wk 10) immunizations were determined using microneutralization assays against the WT SARS-CoV-2 (2019-nCoV/USA-WA1/2020), pseudotyped lentivirus expressing the WT SARS-CoV-2 spike protein (Lenti-CoV2), and MA-SARS-CoV-2. Viral neutralization was plotted as percentage inhibition of viral infection in Vero E6 cells (for WT virus and MA-virus) relative to virus only (no serum) positive controls versus the inverse serum dilution. The titer at which 50% inhibition of infection was achieved (IC50) was determined for the (A, B) WT virus and the (D, E) MA virus. (C) The results were confirmed for the same week 10 serum samples using the Lenti-CoV2 pseudovirus expressing firefly luciferase with HEK-293T cells expressing hACE2. Microneutralization titers using the Lenti-CoV2 were determined by detecting viral infection by measuring luminescence (*p<0.05, **p<0.01 by Mann-Whitney U test). Pretreatment (pre) sera were obtained from the same set of mice before immunizations.

As both WT and MA viruses require the use of BSL-3 containment facilities, to facilitate future vaccine candidate screening, a luciferase-based pseudotyped virus assay was validated. A lentivirus pseudotyped virus expressing the SARS-CoV-2 S protein from the same variant from which the S1 subunit used for immunization was derived (Wuhan-Hu-1), was constructed (Lenti-CoV2), carrying genes for firefly luciferase as described in the methods section. The Lenti-CoV2 S protein contains aa’s 738-1254 of the full length S protein which has a terminal 19 aa deletion removing an ER retention signal. This deletion has been shown to facilitate generation of spike pseudotyped lentivirus^51^.

Microneutralization assays using the WT-SARS-CoV-2 with sera from mice immunized with S1 alone revealed low or undetectable NAb titers, similar to naïve mice after three immunizations (Figure 4A, B). In contrast, mice immunized with NE/S1 showed viral neutralization titers (IC50 GMT 50;range 5-353) after the second immunization (wk 6), which were further increased by two orders of magnitude (IC50 GMT 2.6×10^3^;range 0.4-6.8×10^3^) after the third immunization (wk 10). The combined adjuvant enhanced neutralization titers compared to the NE alone. Sera from mice immunized with NE/IVT/S1 showed increased IC50 values approximately an order of magnitude higher than the NE/S1 group, giving an IC50 GMT of 340 (range 52 to 3.5×10^3^) and IC50 GMT of 8.6×10^3^ (range 4.3×10^3^-3×10^4^) after the second and third immunizations, respectively. Interestingly, this enhancement in virus neutralization was observed with the combined adjuvant, even though there were no differences observed in either the total IgG titers or IgG avidity between the NE/S1 group and the NE/IVT/S1 group at either time point. Notably, sera from NE/IVT/S1immunized mice after the last immunization reached a maximum of ~100% viral neutralization and maintained this level of neutralization even down to serum dilutions often reported as the IC50 for a large proportion of human convalescent sera and for antibodies induced by some lead vaccine candidates in humans and in mice^52–54^. While it is difficult to directly extrapolate results, as neutralization assays still need to be standardized for SARS-CoV-2, these results support induction of high-quality antibodies with the combined adjuvant.

Neutralization titers were confirmed using the Lenti-CoV2 pseudotyped virus in a luciferase based assay with 293T-hACE2 cells (Figure 4C). Microneutralization titers (MNTs) determined by measuring reduction in luminescence (viral infection) with the pseudovirus at week 10 showed almost exact correlation with the traditional microneutralization assay with the WT virus. No NAbs were detectable with this method for the S1 only group, and a similar degree of enhancement in the MNT (~an order of magnitude increase) was observed for the combined adjuvant compared to the NE alone, as was seen in the traditional microneutralization assay.

We next evaluated cross-variant neutralization using the MA-SARS-CoV-2 virus. While slightly reduced compared to titers for WT-SARS-CoV-2 and Lenti-CoV2, high cross-variant neutralization titers were still observed when the sera from NE/IVT/S1 immunized mice were tested against MA-SARS-CoV-2 (Figure 4D, E). There appeared to be a larger reduction in the ability of sera from NE/S1 immunized mice to cross-neutralize the MA-virus, as compared to the NE/IVT/S1 immunized mice. After two immunizations, the difference in enhancement in MNTs in sera from NE/IVT/S1 immunized mice (IC50 GMT 150; range 34-405) compared to NE alone (IC50 GMT 16;range 2.2-360) was greater than the difference observed between the groups with the WT virus due to the larger drop in the ability of the sera from the NE/S1 group to neutralize the MA-virus. However, after the third immunization, differences between the NE and NE/IVT groups became similar to the differences observed with the WT virus, as the MNTs of both groups increased (IC50 GMT 1.2×10^3^; range 420 to 1.9×10^3^ for NE/S1), (IC50 GMT 1.6×10^3^; range 608 to 3.7×10^3^ for NE/IVT/S1). These results further suggest that the combined adjuvant strengthens the quality of the antibody response, providing a protective advantage against divergent variants.

### Passive transfer and challenge

NAb titers required for protection against SARS-CoV-2 have yet to be determined. However, studies in NHPs suggest that low titers (1:50) administered prior to challenge are enough to impart partial protection from a low dose viral challenge, whereas titers of 1:500 conferred full protection to the homologous virus^17^. To determine whether the antibodies raised against the S1 of Wuhan-Hu-1 could protect against heterologous challenge with MA-SARS-CoV-2, week 10 sera from immunized mice were pooled and passively transferred into naïve mice intraperitoneally before challenge IN with 10^4^ PFU virus. While the MA-virus causes mortality and morbidity in aged mice, young C57Bl/6 mice do not lose body weight in this challenge model, as was the case in this study (Figure 5A)^50^. None of the mice receiving serum from immunized animals displayed changes in body weight or increased illness, which also suggests an absence of antibody dependent enhancement (ADE) of disease. Lungs from challenged mice were harvested at 3 d.p.i. for measurement of viral titers in homogenate by plaque assay (Figure 5B). Mice receiving transferred sera from NE/S1 and NE/IVT/S1 groups showed complete sterilizing protection against challenge, with no detectable virus in the lungs. Given the high NAb titers present in both the single and combined adjuvant groups at week 10, it is not surprising that challenge with this moderate viral dose shows no differences between the groups. In contrast, only a slight reduction in viral titers was found for two out of three mice receiving sera from the S1 only immunized mice as compared to the group receiving no serum (PBS control), and one animal in the S1 only group had no viral titers after challenge. We repeated the passive transfer for the PBS control and S1 groups, but this time, virus titers were determined in the whole lung in the repeated challenge. High titers of virus are observed in all of the mice receiving the S1 only sera when whole lung is considered, and again a slight reduction (~10-fold) is observed when compared to whole lung homogenate of mice receiving no serum. This is consistent with the presence of low levels of NAbs in the S1 serum. We are aware of the limitations of this particular MA-SARS-CoV-2 model, as it represents only mild disease in young mice, and repeating the studies with a high dose viral challenge in aged mice would provide greater distinction between the NE and NE/IVT DI groups.

**Figure 5.**
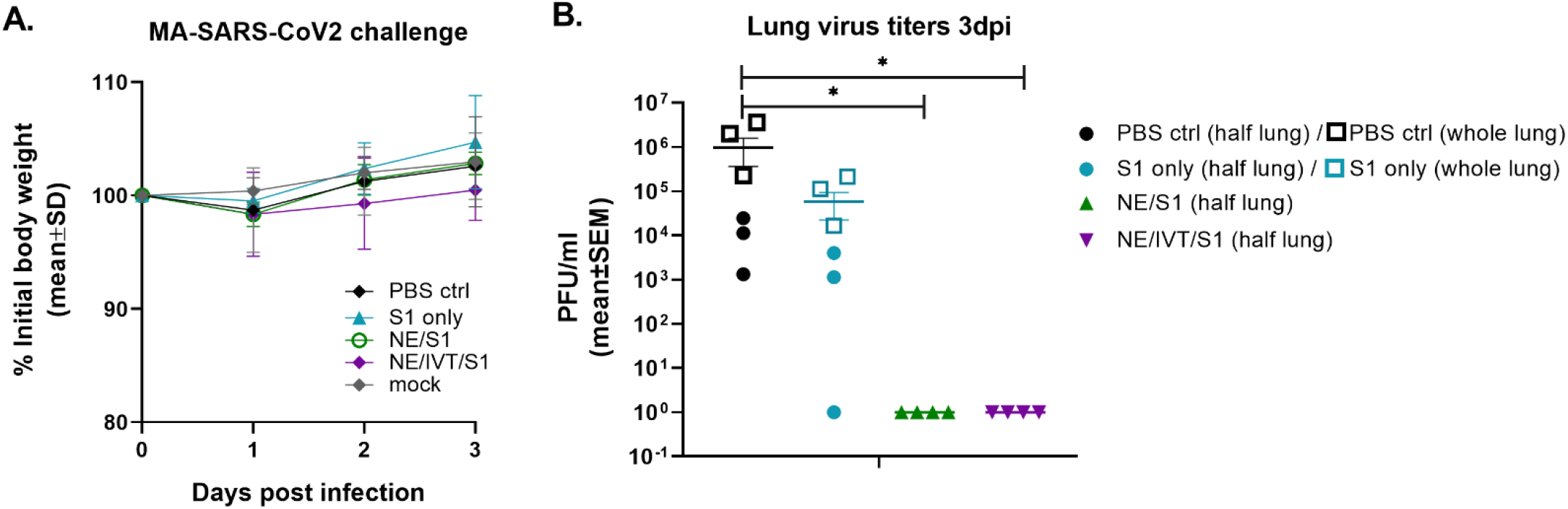
Protection offered by passive transfer of serum from vaccinated mice against heterologous challenge with MA-SARS-CoV-2. Naïve C57Bl/6 mice (n=3-4/group) each received 150 μL of pooled serum through the intraperitoneal route from donor mice given three IN immunizations of S1, NE/S1, or NE/IVT/S1. 2 h after serum transfer, mice were challenged IN with 10^4^ PFU of MA-SARS-CoV-2. (A) Body weight loss was measured over three days, and at 3 dpi (B) lung virus titers were determined in homogenate from one lobe of the isolated lungs by plaque assay (solid symbols). Passive transfer/challenge was repeated for the PBS control and S1 sera for verification, and virus titers were measured in whole lung in the replicated experiment (open square symbols). (*p<0.05, **p<0.01 by Mann-Whitney U test assessed for half lung data points)

### Antigen-specific cellular response profile

Antigen-specific T-cell recall responses were then assessed in splenocytes (Figure 6) and cells isolated from the draining lymph node (cervical lymph node (cLN)) (Figure 7) of the immunized mice two weeks after the last immunization (wk 10). Splenocytes and cLN were stimulated with S1 for 72 h, and cytokine secretion was measured. NE/IVT/S1 administered IN, induced a heavily magnified T_H_1 biased response particularly in the cLN as compared to NE/S1. IFN-γ production in the NE/IVT group was increased by an average of 6-fold and by as high as 60-fold, in the spleen, and increased an average of 10-fold and by as high as 230-fold in the cLN as compared to the NE group (Figures 6A, 7A). IL-2 production in the NE/IVT group was also increased by an average of 2-fold and by as high as 8-fold in the spleen, and increased by an average of 5-fold and by as high as 28-fold in the cLN as compared to the NE group. Additionally, IP-10 and TNF-α were both also enhanced in the spleen and cLN as compared to the NE group. This magnification of T_H_1 associated cytokines and TNF-α is significant, as co-production of IFN-γ, IL-2, and TNF-α on polyfunctional antigen-specific T-cells has been shown to be the strongest criteria for predicting vaccine-elicited T-cell mediated protection against viral infection^55, 56^. Upon analysis of T_H_2 associated cytokines, no significant IL-4 induction was observed in any of the treatment groups, and only minimal levels of IL-13 were observed with NE or NE/IVT that were equivalent to that induced by the antigen alone (Figure 6G,I, 7G,I). NE/IVT immunized mice showed slightly higher levels of IL-5 in splenocytes compared to NE alone, however, levels were low overall, being well below that induced by the S1 alone (Figure 6H). Immunization with NE or NE/IVT actually appeared to reduce the amount of IL-5 and IL-13 induced by the S1 alone (NE/IVT IL-5 was ~5-10-fold lower than the S1 group). While IL-5 was higher in the cLN, a similar pattern applied in which NE and NE/IVT had similar or reduced levels of IL-5 relative to S1 alone (Figure 7H). When IL-5 production after immunization through the IM route with Addavax (MF59) was compared, markedly higher levels of IL-5 (>3,000 pg/mL) were produced in the spleen and cLN upon antigen recall evaluation than with the NE or NE/IVT IN (Figure S1).

**Figure 6:**
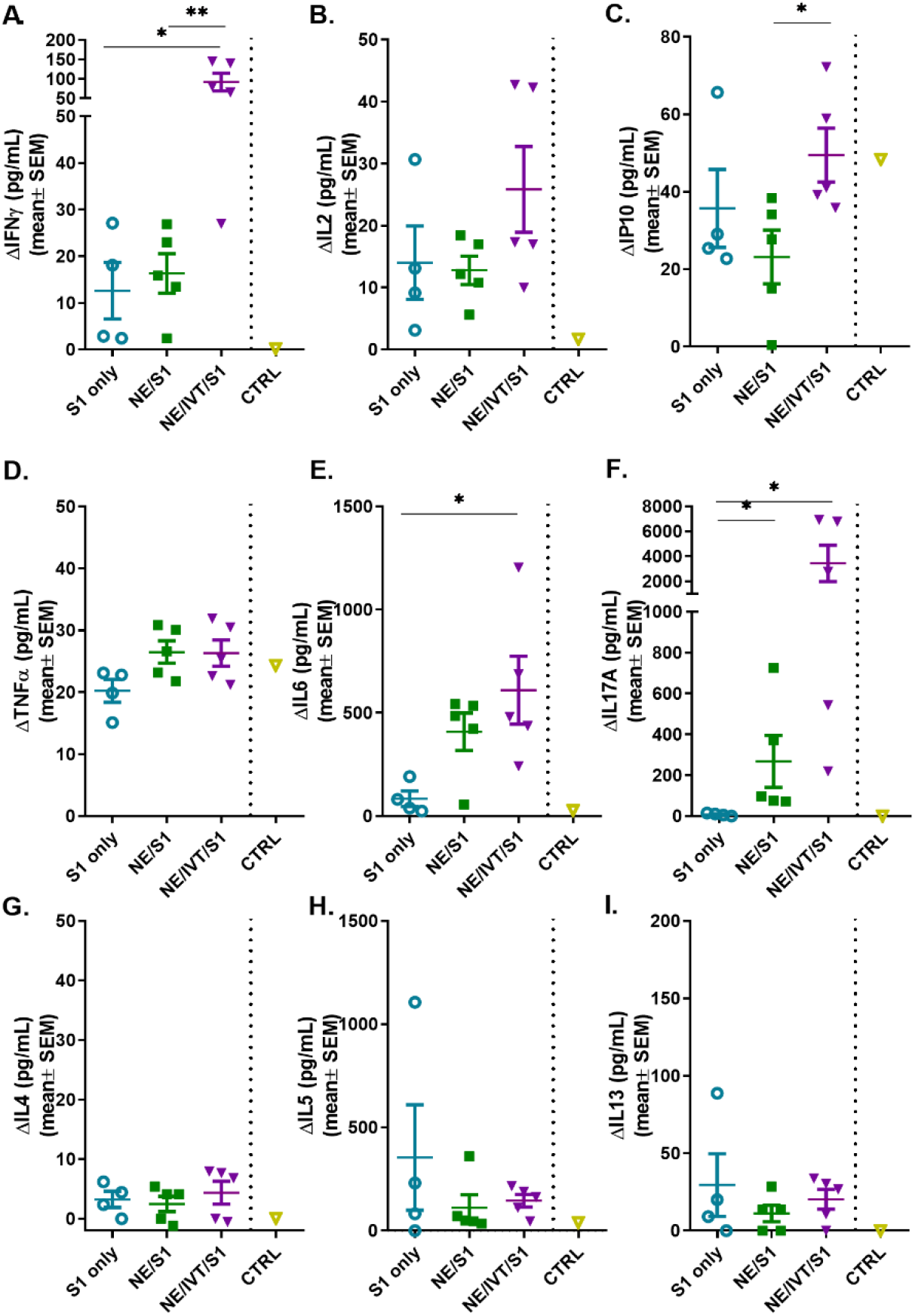
Antigen recall response assessed in splenocytes from immunized animals. Splenocytes isolated from mice immunized IN with S1 alone, or with NE, or NE/IVT after the final boost immunization (10 weeks post-initial immunization) were stimulated *ex vivo* with 5 μg of recombinant S1 for 72 h, and levels of secreted cytokines (A) IFNγ, (B) IL2, (C) IP10, (D) TNFα, (E) IL6, (F) IL17A, (G) IL4, (H) IL5, (I) IL13 were measured in the supernatant relative to unstimulated cells by multiplex immunoassay. An unvaccinated control was included for comparison. (*n*=5/grp; *p<0.05, **p<0.01 by Mann-Whitney U test)

**Figure 7:**
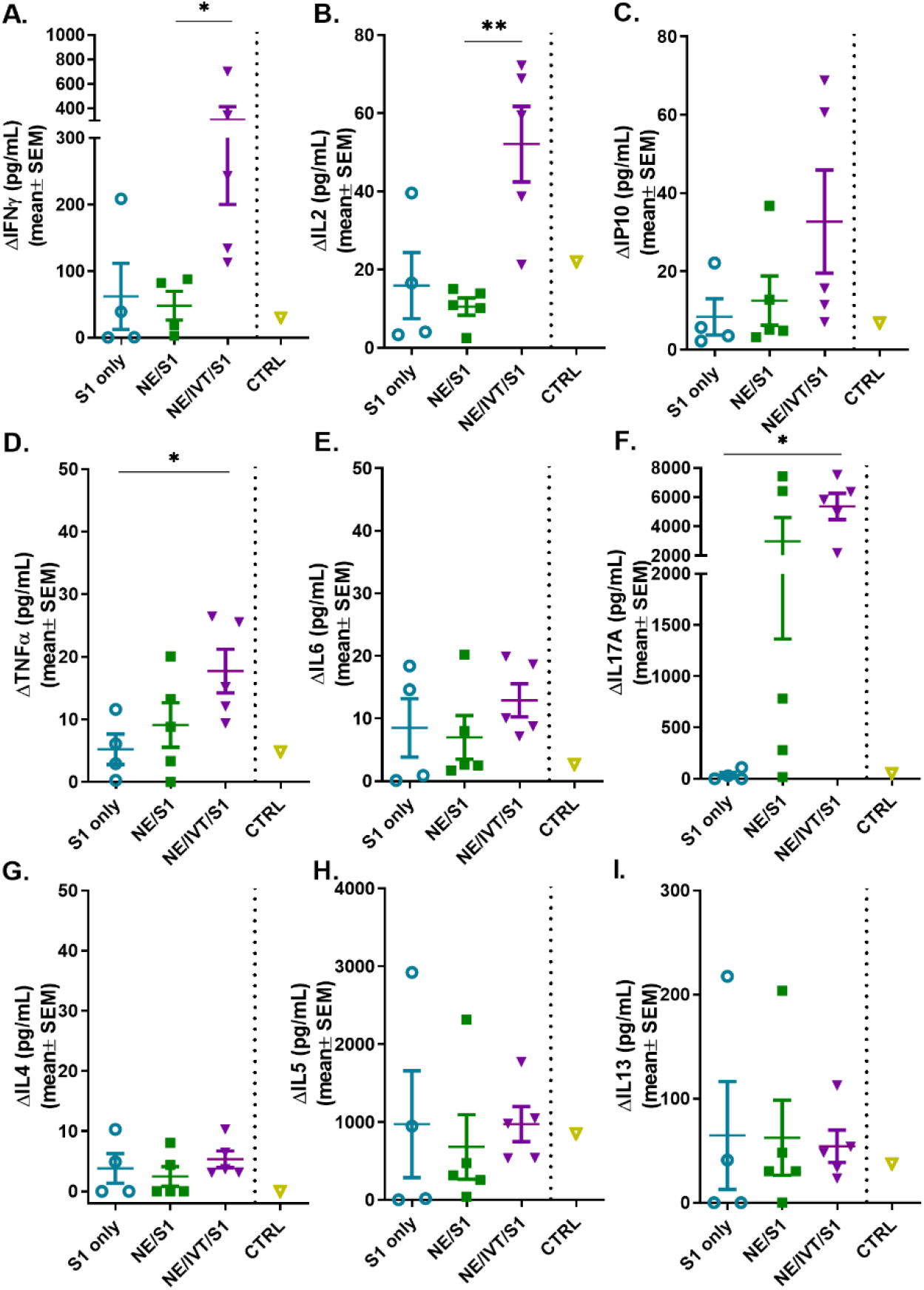
Antigen recall response assessed in the draining lymph nodes (cervical LN). Lymphocytes isolated from the cLNs of mice immunized IN with S1 alone, or with NE, or NE/IVT after the final boost immunization (10 weeks post-initial immunization) were stimulated *ex vivo* with 5 μg of recombinant S1 for 72h, and levels of secreted cytokines (A) IFNγ, (B) IL2, (C) IP10, (D) TNFα, (E) IL6, (F) IL17A, (G) IL4, (H) IL5, (I) IL13 were measured in the supernatant relative to unstimulated cells by multiplex immunoassay. An unvaccinated control was included for comparison. (*n*=5/grp; *p<0.05, **p<0.01 by Mann-Whitney U test)

In addition to the T_H_1 response, a pronounced T_H_17 response as indicated by IL-17A production was also induced by the NE and enhanced significantly by the NE/IVT in the spleen and the cLN. NE/IVT enhanced IL-17A an average of ~10-fold in the spleen, and ~7-fold in the cLN relative to the NE group. We previously observed a similar cytokine response profile including magnified T_H_1 and T_H_17 responses upon immunization of mice with NE/IVT and inactivated influenza virus. Induction of a T_H_17 response is unique to the mucosal route of immunization with NE, and is a critical component of NE-mediated protective immunity through the IN route.

## Discussion

In these studies, we demonstrate the unique efficacy of an intranasal SARS-CoV-2 vaccine containing a rationally designed adjuvant with agonists for all three innate receptor classes important for activating the antiviral response during natural infection. We previously demonstrated that the combination of NE with IVT DI results in an adjuvant that induces synergistic innate responses beyond an additive effect of the individual adjuvants through crosstalk between multivalently activated innate pathways, as well as by NE facilitation of IVT DI cellular uptake. This leads to markedly enhanced production of IFN-Is and improved protective adaptive responses against influenza virus^38^. SARS-CoV-2 activates a dampened innate response characterized by poor IFN-Is induction which leads to highly variable immune responses. In light of the increasing number of variants emerging showing signs of reduced susceptibility to infection- and vaccine-induced immunity, improved strategies to induce effective immune responses which are both potent and broad will be necessary as the virus evolves. Using the NE/IVT adjuvant to provide a more robust and qualitatively appropriate innate cytokine environment strengthened and improved the quality of induced humoral and cellular immune responses to SARS-CoV-2.

Immunization of mice with NE/IVT and S1 resulted in high titers of high avidity antigen-specific IgG after two immunizations, which was further enhanced after an additional boost. NE and NE/IVT elicited high and comparable levels of IgG1 and nearly equivalent IgG2b. For both groups, IgG1 and IgG2b titers were similar, indicating a balanced T_H_1/T_H_2 profile in terms of subclasses. Interestingly, NE/IVT elicited lower IgG2c and slightly reduced IgG2b compared to NE. This is distinct from what we observed in previous studies using NE/IVT with whole inactivated influenza virus, in which the presence of IVT DI (and other RIG-I agonists) enhanced induction of IgG2 subclasses relative to NE alone. These differences may be due to dissimilarities in immune response upon immunization with a whole inactivated virus which contains additional PAMPs, versus a purified protein. Future work will compare the immune responses induced with NE/IVT and inactivated SARS-CoV-2. While the roles of ADCC and ADCP in SARS-CoV-2 immunity are yet unclear, the prevalence of antigen specific IgG2b and 2c is promising for these modes of antibody-mediated immunity.

By designing more effective adjuvants, immune responses directed towards less immunogenic epitopes that are more highly conserved between divergent variants can potentially be better achieved. While NE alone and NE/IVT both yielded similar overall serum antibody titers with similar avidity to S1, NE/IVT induced higher NAb titers to homologous SARS-CoV-2. Moreover, NE/IVT induced antibodies with broader cross-neutralizing capabilities than those generated with NE alone, more effectively neutralizing MA-SARS-CoV-2 which harbors N501Y and H655Y substitutions and a 4aa insertion in S1, supporting the improved quality of the humoral response with the multivalent adjuvant. Passive transfer of antibodies induced after three immunizations with NE and NE/IVT imparted sterilizing immunity in naïve mice against heterologous challenge with MA-SARS-CoV-2. Passive transfer studies performed in NHPs suggest only low NAb titers are necessary for protection against homologous challenge^17^. As the amount of serum transferred was small (150 μl/mouse), it is likely that with such high NAb titers, the number of vaccinations required for inducing sterilizing immunity can be reduced to two or even one with NE/IVT, as we have reported for IM SARS-CoV-2 vaccination in mice with a TLR7/8 agonist as adjuvant which induced significantly lower NAb titers^57^. The ability of NE/IVT to improve cross-variant protection induced with just the S1 subunit, particularly towards a variant with the N501Y mutation shared by several of the highly transmissible variants of concern, supports its potential for improving the breadth of protection of current vaccine candidates. Furthermore, as NE/IVT is compatible with both recombinant protein-based antigens and whole inactivated viruses, utilizing the adjuvant with strategically designed antigens (ex. chimeric antigens) and/or heterologous prime-boost antigen regimens as has been done for influenza vaccines could better hone responses towards regions which are more highly conserved between drift variants^58, 59^.

The importance of cellular immunity in complete protection against SARS-CoV-2 has become clear, with strong correlations found between disease severity and the presence of CD4^+^ and CD8^+^ T cell responses. Memory CD8^+^ T cells, especially tissue resident memory T cells (T_RM_’s) have been shown to be essential to viral clearance of SARS-CoV^60–63^. Moreover, CD8^+^ T cells were longer lived in SARS-CoV convalescent patients than CD4^+^T and B cells, providing protection in their absence^64^. The quality and durability of neutralizing antibody responses and B cell memory depend tightly upon CD4^+^ T cell help, and CD4^+^ T cells are involved in a wide spectrum of activities critical to antiviral immunity^65^. NE/IVT strongly enhanced cellular immune responses relative to the NE alone, polarizing responses towards a strong T_H_1 bias, with significantly magnified IFN-γ, and increased IL-2, IP-10 and TNF-α production upon antigen recall in the spleen and cLN. In contrast, T_H_2 associated cytokines, IL-4, IL-5, and IL-13 were not increased compared to immunization with S1 alone. VAERD reported for some SARS-CoV vaccine candidates was primarily attributed to poor antibody quality in the context of adjuvant (alum) enhanced T_H_2 immunopathology. In these cases, accentuated IL-4, IL-5, and IL-13 production in vaccinated animals upon viral exposure resulted in potentiation of airway inflammation, hyperresponsiveness, and lung dysfunction, particularly in aged animals^46, 47^. NE/IVT induces a cytokine environment heavily favoring T_H_1 responses with potent T cell activation and high-quality antibody responses, which provides an optimal profile for avoiding VAERD and ADE.

Induction of a T_H_17 response is unique to the mucosal route of NE immunization, and is an important component of NE-mediated protective immunity. While the role of T_H_17 responses in SARS-CoV-2 immunity is not clear, T_H_17 responses promote effective immunity to several respiratory viruses, including influenza virus, in which it enhances viral clearance and survival^66,67^. It is likely that inducing IL-17A *early*, in the context of an appropriate cytokine milieu, will be important for driving protective immunity to SARS-CoV-2. T_H_17 lineage CD4^+^ T cells drive mucosal immunity which is especially important for respiratory viruses as it can prevent transmission and viral dissemination to the lung. Indeed, NE and NE/IVT induced significant antigen-specific IgA in the bronchial alveolar lavage (BAL) of immunized animals (Figure S2). T_H_17 CD4^+^ T cells are also critical for development of T_RM_’s residing in the lung mucosa, which are critical elements of local immunity to SARS-CoV-2, functioning early before circulating effector and central memory cells are recruited^68, 69^. While IL-17A has been associated with lung pathology in certain situations, this is primarily observed in the context of high T_H_2 cytokines, whereas pathological inflammatory effects of IL-17A are prevented by IL10^70^. While IL10 was not measured in the current study, our previous work with other antigens including SARS-CoV-2 RBD (data not shown) and influenza virus among others, have demonstrated significant enhancement of IL10 with NE and NE/IVT^30, 38, 71, 72^. As NE and NE/IVT do not induce high levels of T_H_2 cytokines, while enhancing T_H_1 cytokines and IL-10, this T-cell activation profile will likely lead to enhanced viral clearance without detrimental inflammation, as we have observed in our prior influenza virus challenge studies. IL-17A induced by NE vaccination improved protection to RSV without increased lung pathology or eosinophilia upon challenge in mouse and cotton rat models, further supporting this^73^.

The COVID-19 pandemic is a quickly evolving landscape, with much still being learned regarding the correlates of protection to SARS-CoV-2. Together, our data demonstrate that a combined adjuvant approach offers a promising strategy for promoting both robust and broader antibody and T cell responses to improve protection against SARS-CoV-2. Furthermore, the ability to use NE/IVT DI through the IN route offers the advantages of inducing mucosal immunity in the respiratory tract, the natural route of viral entry. This formulation is amenable to needle-free administration, and is inexpensive and rapidly scalable, making it ideal for mass vaccinations. As NE/IVT DI is compatible with both whole viral antigens and recombinant antigens, it thus provides a flexible platform that has the potential to improve the immune profiles of multiple vaccine candidates currently in development for SARS-CoV-2.

## Materials and Methods

### Adjuvants and antigen

NE was produced by emulsification of cetylpyridinium chloride (CPC) and Tween 80 surfactants, ethanol (200 proof), super refined soybean oil (Croda) and purified water using a high speed homogenizer as previously described^29^. CPC and Tween80 were mixed at a 1:6 (w/w) ratio, and homogeneity of particle size (*d=*450-550 nm) and charge (zeta potential=50-55mV) were confirmed. Stability was assessed over several months. Sequence and synthesis methods for IVT DI RNA have previously been reported in detail^36^. Briefly, SeV DI RNA from SeV-infected A549 cells was amplified using a 5’ primer with the T7 promoter and a 3’ primer with the hepatitis delta virus genomic ribozyme site followed by the T7 terminator. The resultant DNA was cloned into a pUC19 plasmid and *in vitro* transcribed using a HiScribe T7 High Yield RNA synthesis kit (New England Biolabs). After DNAseI digestion and clean-up with a TURBO DNA-free kit (Thermo-Fisher), IVT DI was purified using an RNeasy purification kit (Qiagen). The absence of endotoxin was verified by limulus amoebocyte lysate assay. Recombinant SARS-CoV-2 spike protein S1 subunit (Wuhan-Hu-1 (Val16-Arg685) (accession YP_009724390.1)) with a C-terminal His tag was purchased from Sino Biological.

### Cell lines

Vero E6 cells (ATCC) were maintained in MEM supplemented with 10% heat inactivated fetal bovine serum (HI FBS). HEK293T cells expressing hACE2 (293T-hACE2) were obtained from BEI resources and maintained in HEK293T medium: DMEM containing 4 mM L-glutamine, 4500 mg/L L-glucose, 1 mM sodium pyruvate and 1500 mg/L sodium bicarbonate, supplemented with 10% HI FBS as previously described^74^.

### Viruses

WT SARS-CoV-2: SARS-CoV-2 clinical isolate USA-WA1/2020 (BEI resources; NR-52281), referred to as the WT virus herein, was propagated by culture in Vero E6 cells as previously described^75^. MA SARS-CoV-2: Mouse-adapted SARS-CoV-2 was obtained by serial passage of the USA-WA1/2020 clinical isolate in mice of different backgrounds over eleven passages, as well as on mACE2 expressing Vero E6 cells as previously described^50^. Briefly, the virus was passaged every two days via IN inoculation with lung homogenate derived supernatants from infected mice. All viral stocks were analyzed by deep sequencing to verify integrity of the original viral genome.

### Lentivirus pseudotyped virus

Cloning of expression constructs: For generation of spike protein pseudotyped lentivirus (Lenti-CoV2), a codon optimized SARS-CoV-2 spike protein (accession #QHD43416.1) construct was obtained from Sino Biologicals. All cloning and lentivirus production was performed by the University of Michigan Vector Core. The SARS-CoV-2 spike with 19 amino acids deleted (SΔ19) was generated by PCR amplifying the region of spike containing amino acids 738 to 1254 from the full length SARS-CoV-2 spike construct using following primers; Spike BsrGI Gib Fwd 5’-GACCAAGACCTCTGTGGACTGTACAATGTATATCTGTGGAGAC and Spike Δ19 XbaI Rev 5’-GCCCGAATTCGGCGGCCGCTCTAGAGTTCAACAACAGGAGCCACAGGAAC. The product was cloned into a pCMV3 vector digested at the BsrGI/XhoI sites. Resulting clones were verified by Sanger sequencing. The resulting clone was designated pCMV3-SΔ19. For the generation of a lentiviral vector containing SARS-CoV2–SpikeΔ19, the pCMV3-SΔ19 insert was initially digested with KpnI and blunt polished using Phusion Taq polymerase followed by a DNA cleanup using the Monarch PCR cleanup kit (NEB) and a second digest was done using NotI. The released fragment was then ligated into a pLentiLox-RSV-CMV-Puro vector. Correct insertion was verified by Sanger sequencing. The resulting clone was designated pSARsCoV2Δ19AA.

To prepare Lenti-CoV-2 pseudovirus expressing the SARS-CoV-2 S protein, lentivirus packaging vectors psPAX2 (35 μg), and coronavirus truncated spike envelope pSARsCoV2Δ19AA (35 μg) were co-transfected with 70 μg of pGF1-CMV proviral plasmid using standard PEI precipitation methods. PEI precipitation was performed by incubating the plasmids with 420 μg PEI (molecular weight 25,000, Polysciences, Inc) in 10 mL Opti-MEM (Life technologies) at room temp for 20 m, before adding to fresh 90 mL of DMEM media supplemented with 10% FBS-1XGlutamax-100U/mL Penn/Strep. This DNA/PEI containing media was then distributed equally to 5-T150 flasks (Falcon) containing 293T cells. Supernatants were collected and pooled after 72 h, filtered through a 0.45 μm GP Express filter flask (Millipore), pelleted by centrifugation at 13,000 rpm on a Beckman Avanti J-E centrifuge at 4°C for 4 h, and resuspended at 100X the original concentration (~1×10^7^ TU/ml) in DMEM. Harvested lentivirus was stored at −80 C.

### Animals

All animal procedures were approved by the Institutional Animal Care and Use Committees (IACUC) at the University of Michigan and Icahn School of Medicine at Mt. Sinai and were carried out in accordance with these guidelines. 6-8-wk-old female C57Bl/6 mice (Jackson Laboratory) were housed in specific pathogen-free conditions. Mice were acclimated for 2 wks prior to initiation. For challenge studies, mice were transferred to ABSL3 facilities 2 d prior to serum transfer and subsequent viral challenge.

### Immunization

For intranasal (IN) immunization, mice were anesthetized under isoflurane using an IMPAC6 precision vaporizer and given 12 μL total (6 μL/nare) of each vaccination mixture. Each group received a total of three immunizations of the same formulations at 4-wk intervals. 15 μg of S1 was administered with either PBS, 20% NE (w/v), or 20% NE with 0.5 μg of IVT DI in PBS. Sera were obtained by saphenous vein bleeding 2 and 4wks after each immunization, and by cardiac puncture at the end of the experiment at week 10. Bronchial alveolar lavage (BAL) was obtained by lung lavage with 0.8 mL PBS containing protease inhibitors. Spleens and cervical lymph nodes were harvested, processed to single-cell suspensions, and cultured for antigen recall response assessment as previously described^38^.

### ELISA

Immunograde 96-well ELISA plates (Midsci) were coated with 100 ng S1 in 50 μL PBS per well overnight at 4°C, and then blocked in 200 μL of 5% non-fat dry milk/PBS for 1 h at 37°C. Serum samples from immunized mice were serially diluted in PBS/0.1% BSA starting at either a dilution of 1:50 or 1:100. Blocking buffer was removed, and diluted sera were added to the wells and incubated for 2 h at 37°C followed by overnight incubation at 4°C. Plates were washed three times with PBST (0.05% Tween20), and alkaline phosphatase conjugated secondary antibodies diluted in PBS/0.1% BSA were added (goat-anti-mouse IgG, IgG1, IgG2b, or IgG2c Jackson Immuno Research Laboratories). Plates were incubated at 37°C for 1h, washed with PBST, and then developed at RT by addition of 100 μL of p-nitrophenyl phosphate (pNPP) substrate (Sigma-Aldrich) per well. Absorbance was measured at 405nm on a microplate spectrophotometer. Titers were calculated against naïve sera, using a cutoff value defined by the sum of the average absorbance at the lowest dilution and two times the standard deviation.

### Pseudovirus microneutralization (MNT) assays

9×10^3^ 293T-hACE2 cells were seeded overnight on white clear bottom 96-well tissue culture plates in HEK293T medium. To titer the virus, the Lenti-Cov2 stock was serially diluted in HEK293T medium with 16μg/mL polybrene (Sigma-Aldrich), and incubated for 1h at 37°C to mimic assay conditions for MNT. Diluted virus was then added to the 293T-hACE2 cells and incubated at 37°C for 4 h. The media was replaced with fresh HEK293T medium without polybrene and incubated for an additional 72 h at 37°C. Infection medium was removed and replaced with 20 μL of BrightGlo luminescence reagent using an injection luminometer. Cells were incubated for 2 m with shaking and luminescence was collected over a read time of 1 s. For MNT, 293T-hACE2 cells were seeded overnight. Serum samples from immunized mice were serially diluted by a factor of two, starting at a dilution of 1:50 in HEK293T medium with 16 μg/mL polybrene (Sigma-Aldrich). 50 μL of diluted sera was added to 50 μL of lenti-CoV2 in the same media at a concentration which gave a luminescence reading of ≥20,000 RLU/well above background in infected cells as determined by viral titration. Serum and virus were incubated for 1h at 37°C, and then added to 293T-hACE2 cells for incubation at 37°C for 4 h. Infection medium was removed, and replaced with fresh HEK293T medium without polybrene and incubated for an additional 72 h at 37°C. Luminescence was measured as above. Neutralization titers were determined as the dilution at which the luminescence remained below the luminescence of the (virus only control-uninfected control)/2.

### Microneutralization Assays

MNT assays with WT SARS-CoV-2 (2019-nCoV/USA-WA1/2020) and the mouse adapted (MA-SARS-CoV-2) variant was performed in a BSL3 facility as previously described^50^. Briefly, 2×10^4^ Vero E6 cells were seeded per well in a 96-well tissue culture plate overnight. Serum samples were heat-inactivated for 30 m at 56°C and serially diluted by a factor of 3, starting at dilutions of 1:10 or 1:20 in infection medium (DMEM, 2% FBS, 1x non-essential amino acids). Diluted serum samples were incubated with 450xTCID50 of each virus which (~40 PFU) for 1 h at 37°C. Growth medium was removed from the Vero E6 cells, and the virus/serum mixture was added to the cells. Plates were incubated at 37°C for 48 h, fixed in 4% formaldehyde, washed with PBS and blocked in PBST (0.1% Tween 20) for 1 h at RT. Cells were permeabilized with 0.1% TritonX100, washed and incubated with anti-SARS-CoV-2-nucleoprotein and anti-SARS-CoV-2-Spike monoclonal antibodies, mixed in 1:1 ratio, for 1.5 h at RT. After another wash, cells were incubated with HRP-conjugated goat-anti-mouse IgG secondary antibody for 1h at RT. Cells were washed, and plates were developed by incubation with 50 μL tetramethyl benzidine until a visible blue color appeared, after which the reaction was quenched by addition of 50 μL 1M H_2_SO_4_. Absorbance was measured at 450nm and percentage inhibition (reduction of infection) was calculated against virus only infected controls. The 50% inhibitory dilution (IC50) values were calculated for each sample. Undetectable neutralization was designated as a titer of 10^0^. Anti-mouse SARS-CoV-2-nucleoprotein and anti-mouse SARS-CoV-2-spike antibodies were obtained from the Center for Therapeutic Antibody Development at the Icahn School of Medicine at Mount Sinai.

### Antigen Recall Response

T cell antigen recall response was assessed in cell isolates from the spleen and cLN of immunized mice 2wks after the final immunization (week 10). Methods for splenocyte and cLN lymphocyte preparation were previously described^38^. For antigen recall, isolated cells were plated at a density of 8×10^5^cells/well and stimulated with 5 μg per well of recombinant S1 in T cell media (DMEM, 5% FBS, 2 mM L-glutamine, 1% NEAA, 1 mM sodium pyruvate, 10 mM MOPS, 50 μM 2-mercaptoethanol, 100 IU penicillin, and 100 μg/mL streptomycin), in a total volume of 200 μL per well. Cells were stimulated for 72h at 37°C, and secreted cytokines (IFN-γ, IL-2, IL-4, IL-5, IL-6, IL-17A, TNF-α, and IP-10) were measured relative to unstimulated cells in supernatants using a Milliplex MAP Magnetic Mouse Cytokine/Chemokine multiplex immunoassay (EMD Millipore).

### Acute cytokine response

Acute response markers were measured in immunized mice 6 and 12 h post-initial immunization to determine whether the formulations induced reactogenicity. For each group, mice were bled either at 6 or 12 h and the amounts of IFNγ, IL6, IL12p70, and TNFα in the sera were measured using a Procartaplex multiplex immunoassay (ThermoFisher) according to the manufacturer’s protocol.

### Passive Transfer and Challenge

Serum samples were pooled from mice in each immunization group collected after the second boost immunization (wk 10), and 150 μL of the pooled serum was passively transferred into naïve mice through the intraperitoneal route 2 h prior to challenge intranasally under mild ketamine/xylazine sedation with 10^4^ PFU of MA-SARS-CoV-2 in 30 μL. Body weight changes were recorded every 24h, and mice were sacrificed at 3 d.p.i. Lungs were harvested and homogenate prepared for virus titration by plaque assay as previously described^50^.

### Avidity

To measure antibody avidity of serum IgG in immunized mice, ELISAs were performed as described above against S1 modified by an additional chaotrope elution step after the overnight incubation of serum on the ELISA plate as previously described^38^. Briefly, after the serum incubation and washes with PBST, 100 μL of PBS, or 0.5 M NaSCN, or 1.5 M NaSCN in PBS at pH 6 was added to each well and incubated at RT for 15 min. The plates were washed three times with PBST, and the ELISA proceeded to development as described above by addition of alkaline phosphatase conjugated goat-anti-mouse IgG.

## Acknowledgements

We thank the U of M vector core and Dr. Tom Lanigan for producing the lentivirus pseudovirus and for providing technical input on assays. The authors gratefully thank Adolfo García-Sastre and Raffael Nachbagauer at the Icahn School of Medicine at Mount Sinai, New York, NY for support, use of laboratory infrastructure and helpful discussions. We also thank Richard Cadagan for excellent technical assistance and Randy Albrecht for BSL3 support. This research was partly funded by CRIP (Center for Research for Influenza Pathogenesis), a NIAID supported Center of Excellence for Influenza Research and Surveillance (CEIRS, contract # HHSN272201400008C), by supplements to NIAID grant U01AI124297 and by Collaborative Influenza Vaccine Innovation Centers (CIVIC) contract 75N93019C00051.

## Supplemental Information

**Supplemental Figure S1:**
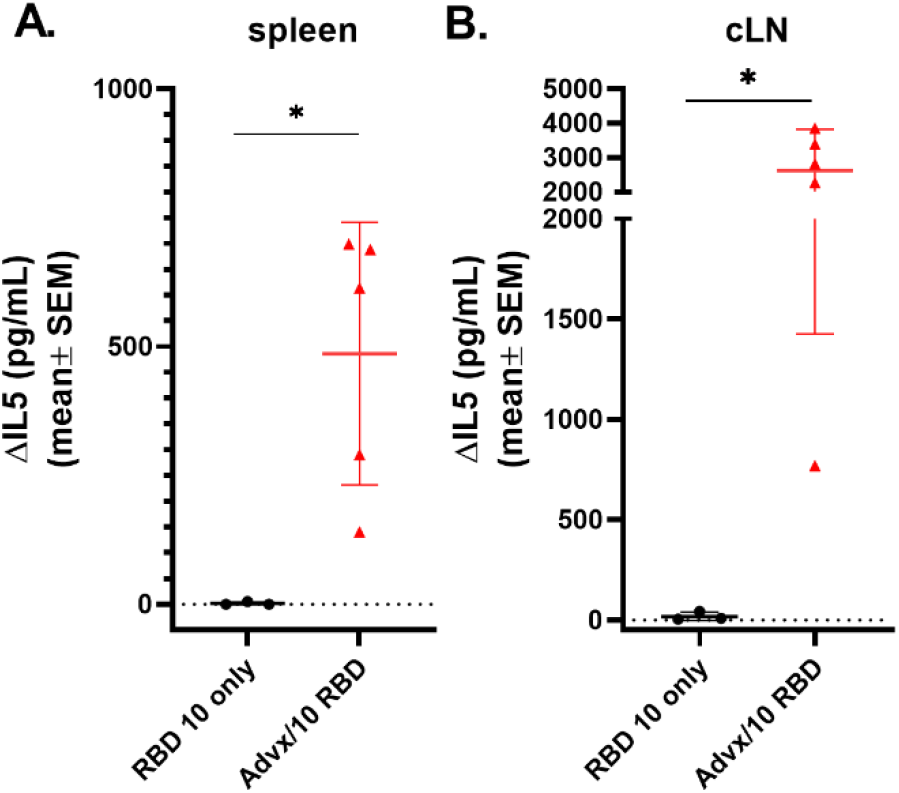
Antigen recall response assessed in lymphocytes from spleen and cLN isolated from mice immunized IM with 10 μg SARS-CoV-2 RBD alone, or with 50% Addavax in a volume of 50 μL according to a prime/boost/boost schedule (at a 4 wk interval). Cells were stimulated *ex vivo* with 5 μg of recombinant RBD for 72h, and levels of secreted IL-5 were measured in the supernatant relative to unstimulated cells by multiplex immunoassay. (*n*=5/grp; *p<0.05, **p<0.01 by Mann-Whitney U test)

**Supplemental Figure S2.**
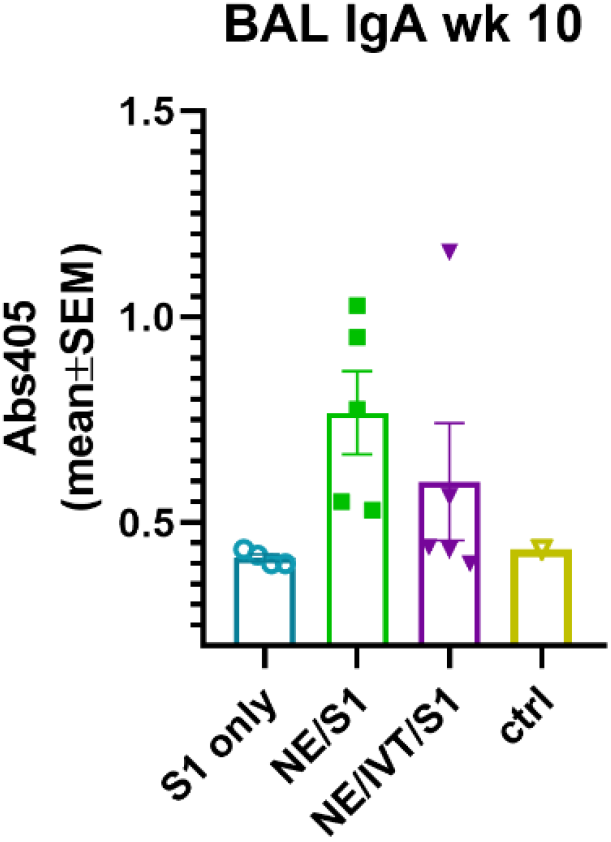
Mucosal response assessed in immunized mice by measuring S1-specific IgA in bronchial alveolar lavage after prime/boost/boost immunizations (10 weeks post-initial immunization) as measured by ELISA. Absorbance values at 405 nm are shown after development with an alkaline-phosphatase conjugated secondary antibody with a pNPP substrate.

